# Illuminating dark genes and dark metabolites: calibrated functional association across kingdoms with autoencoders and correlation

**DOI:** 10.64898/2026.03.26.712966

**Authors:** Yannick Mahlich, Dylan H. Ross, Lummy M. O. Monteiro, Sutanay Choudhury, William C. Nelson, Jason E. McDermott

## Abstract

**Motivation:** Despite continuing advances in sequencing and computational function prediction, large parts of gene, protein, and metabolite space remain functionally undetermined. Most function assignment is based on homology searches and annotation transfer from well-studied proteins, but this approach fails to leverage available experimental omics data generated via technologies like mass-spectrometry.

**Results:** To address this gap, we have developed the VaLPAS (Variation-Leveraged Phenomic Association Screen) framework, a Python package that provides a user-friendly platform to discover associations between expression patterns of genes, proteins, and metabolites in multi-omic datasets. VaLPAS uses various statistical and learning methods, from simple statistical methods (correlation) to a sophisticated autoencoder, to shed light on the functional dark matter of protein space using ‘guilt by association’ with molecules of known function. We have used VaLPAS to examine transcript, protein, and metabolite data from seven organisms across two kingdoms using two different benchmarks. We present results demonstrating the ability of VaLPAS to make high-confidence predictions of the functional associations of a subset of genes and proteins. Furthermore, we show that VaLPAS can provide insight into the functional relevance of detected but unidentified metabolites by cross-omics association, a unique feature of this approach. This ability of cross-omics association enables VaLPAS to serve as a tool for generating hypotheses of functional interaction that can be validated with targeted experiments, representing a significant step forward in elucidating dark functional matter.

**Availability and implementation:** VaLPAS is hosted on github (https://github.com/PNNL-Predictive-Phenomics/valpas/).

**Contact:** Jason E. McDermott

**Supplementary information:** Supplementary information available online at Oxford Bioinformatics

## 1. Introduction

Genetic sequencing data deposited in public databases continues to grow at an ever-increasing pace (Consortium, 2024; O’Leary, et al., 2016). Yet experimental verification of functional predictions for gene products has not remotely kept pace. The vast majority of protein sequences in public databases have been annotated using sequence similarity to infer homology relationships and transferring function through these associations. Even in well-established model organisms such as *Escherichia coli* and *Saccharomyces cerevisiae*, between 20% and 35% of protein-coding genes remain poorly characterized or entirely lack functional annotation (Lobb, et al., 2020; Salzberg, 2019). While recent advances in protein structure prediction (Ahdritz, et al., 2024; Jumper, et al., 2021; Mirdita, et al., 2022) are enabling new methods leveraging structural similarity to improve function prediction (Gligorijevic, et al., 2021; Wang, et al., 2025), these methods still suffer from the same problems as function transfer by sequence similarity, i.e. they have significantly reduced predictive power if evolutionary relationships are unclear (Kryshtafovych, et al., 2023; Ozden, et al., 2023). Initiatives like CAFA (Critical Assessment of Functional Annotation) have been a driving factor in advancing the field of protein function prediction in the last decade, yet still face the same challenges outlined above (Jiang, et al., 2016; Piovesan, et al., 2024; Radivojac, et al., 2013; Zhou, et al., 2019).

The discussion of functional characterization of molecular components in biological systems is often restricted to genes and proteins, whereas small molecule metabolites are neglected. Despite technical progress typically allowing hundreds of metabolites to be consistently identified and measured using modern untargeted mass spectrometry approaches, it is estimated that over 95% of measured peaks corresponding to potential molecular species in metabolomics experiments can’t be assigned a molecular structure (da Silva, et al., 2015). This prevents the use of a majority of metabolomics data and limits its utility in helping understand system function. Additionally, as with proteins, metabolites can be associated with more than one metabolic pathway. Current understanding of their roles in bacterial and fungal systems is captured in canonical pathway definitions presented in databases like the Kyoto Encyclopedia of Genes and Genomes (KEGG) (Kanehisa, et al., 2025).

The growing field of biological AI attempts to address this challenge of functional prediction without the reliance on homology as a proxy of evolutionary information. Specialized language models for biological context such as protein language models (pLMs) as well as more complex learning structures such as graph neural networks incorporating additional prior knowledge hold promise for better prediction of function (Rives, et al., 2021; Yu, et al., 2023), though limitations still exist for many such methods that use sequence as their only data source (Olson, et al., 2025; Prabakaran and Bromberg, 2026).

The set of methods collectively known as ‘guilt-by-association’ uses patterns encoded in omics abundance patterns, commonly co-expression, to associate genes from the same experiment and provide insight for unannotated features (Wang, et al., 2017; Wolfe, et al., 2005). In the simplest approaches demonstrated to be effective, statistical correlation between molecules across different conditions has been used (Wolfe, et al., 2005). Deep learning approaches like autoencoders offer the potential for capturing non-linear patterns in omics datasets and potentially outperforming statistical methods (Way, et al., 2020). However, no systematic evaluation of when each approach is preferable across organisms, omics modalities, and sample sizes using easily understandable confidence measures has been undertaken. Furthermore, the potential for these approaches to provide cross-omics predictions, i.e. predicting function of unknown metabolites by their association with known protein pathways, has not been assessed.

To this end we introduce VaLPAS (Variation-Leveraged Phenomic Association Screen) a Python framework that enables researchers to easily find associations between different matched omics data points from high-throughput studies such as transcriptomics, proteomics, and metabolomics. VaLPAS is a general framework that provides capability for association studies from multi-omics data using correlation, distance metrics as well as a novel bi-directional autoencoder to learn non-linear patterns from omics experiments. VaLPAS serves as a tool to uncover and systematically explore novel molecular associations, providing additional functional information for genes, proteins and metabolites not captured by sequence similarity methods.

We compare the performance of VaLPAS’ correlation and autoencoder-based guilt-by-association analysis of multiple datasets across bacterial and fungal species, with transcriptomics, proteomics, and metabolomics, establishing a confidence baseline for further evaluations. Using these baselines, we establish functional associations of dark genes in ***Pseudomonas putida*** and compare those with results from a recent study by Tavis and Hettich (Tavis and Hettich, 2024), revealing high confidence, yet orthogonal predictions. Additionally, we perform an analysis investigating dark metabolites in ***Lipomyces starkeyi*** based on a recently published study by Monteiro et al. (Monteiro, et al., 2026). In both cases we successfully identify high-confidence functional associations for unannotated genes and metabolites with known and functionally annotated proteins. Simultaneously, these analyses reveal that data-dependent differences determine which approach provides better predictions.

## 2. Methods

### 2.1 Datasets

#### 2.1.1 Bacterial and fungal (multi-)omics data

To fully evaluate the capabilities of VaLPAS and to compare the statistical approaches with autoencoder, 7 existing published datasets (4 bacterial, 3 eukaryotic) consisting of different sets of omics data were obtained (see Table 1). Datasets with multiple omics data types were matched by sample. Bacterial transcriptomics datasets (*Escherichia coli*, *Pseudomonas putida*, *Salmonella enterica*, *S. elongatus*) were downloaded from the iModulon database (Catoiu, et al., 2025). *Saccharomyces cerevisiae* transcriptomics data was downloaded from GEO (GSE42527) (Kemmeren, et al., 2014). Multi-omics for *Rhodotorula toruloides* (transcriptomics, proteomics and metabolomics) and *Lipomyces starkeyi* (proteomics and metabolomics) were retrieved from Kim, et al. 2020 (Kim, et al., 2020) and Monteiro, et al. 2026 (Monteiro, et al., 2026) respectively.

**Table 1.** Datasets used in this study (T, transcriptomics; P, proteomics; M, metabolomics)

| Organism | Domain | Omics | Samples | Conditions |
| --- | --- | --- | --- | --- |
| *E. coli* K-12 | Bacteria | T | 3,376 | Compendium |
| *P. putida* KT2440 | Bacteria | T | 321 | Various growth |
| *S. enterica* LT2 | Bacteria | T | 533 | Various growth |
| *S. elongatus* | Bacteria | T | 139 | Various growth |
| *S. cerevisiae* S288C | Eukarya | T | 700 | Stress/nutrients |
| *R. toruloides* NP11 | Eukarya | T,P,M | 11 | N-limitation time |
| *L. starkeyi* Y11558 | Eukarya | P,M | 36 | C×N×treatment |

#### 2.1.2 Data processing

Input data is organized as measurement matrices where rows represent biological conditions (samples) and columns represent omics features (genes, proteins, metabolites, etc.). VaLPAS supports single-omics analysis (one measurement matrix) and cross-omics analysis (two measurement matrices from different omics types sharing the same conditions). The input data is individually preprocessed on a per analysis and per dataset basis, to filter out omics features with high missingness, i.e. features that have not been detected in enough sample conditions (user defined; default: missingness <= 10%, i.e. the feature has to be present in at least 90% samples). Optional normalization can be specified prior to processing using Z-score normalization, pareto, or power scaling.

### 2.2 VaLPAS

VaLPAS allows users to process their omics abundance data matrices to retrieve associations between pairs of molecular features indicated by scores specific to the association method chosen (Figure 1). Those include correlation (Pearson and Spearman), mutual information using bsplines (Faith, et al., 2007), cosine and Jaccard similarity. For these association methods, scores are calculated based on pairs of abundance vectors which record the features’ abundances across all measured samples. Furthermore, VaLPAS includes an additional bi-directional autoencoder architecture that learns associations through self-supervised masked learning (see Section 2.3). Notably, the calculation of association scores using the autoencoder architecture differs from all other methods in that the autoencoder considers not only the abundances of the compared features across all samples, but also encodes samples based on the feature abundances per sample. Returned association scores for features extracted using the autoencoder architecture are cosine similarities between feature embedding vectors. Note this is different from the previously mentioned non-autoencoder architecture based cosine similarity.

**Figure 1.**
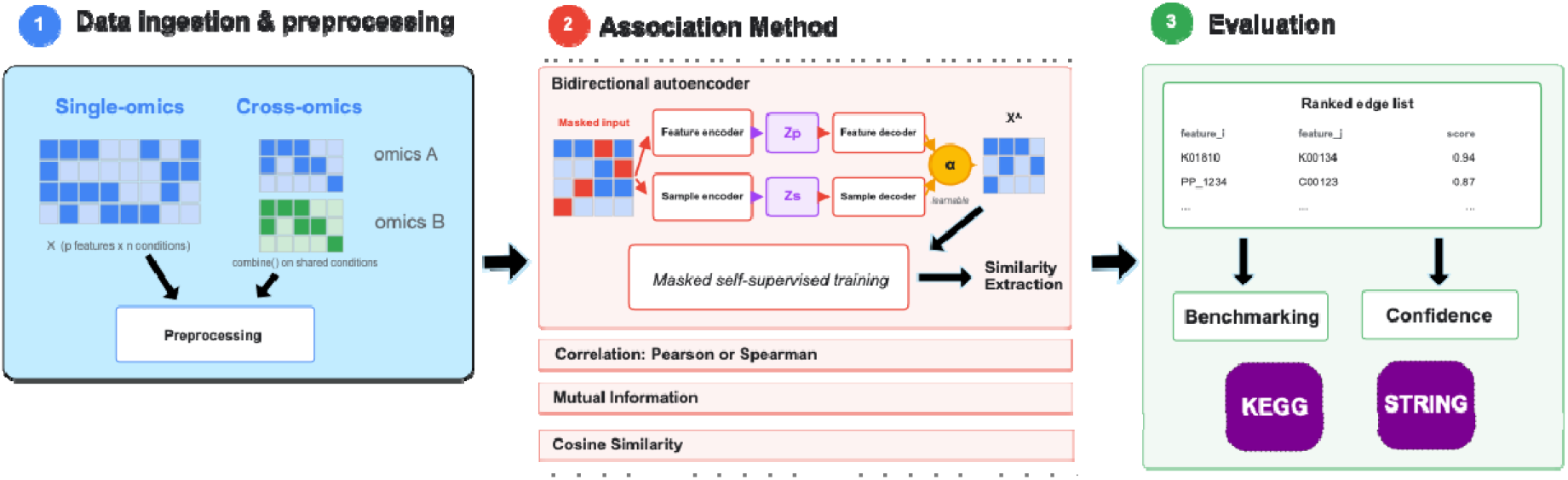
VaLPAS workflow and architecture. Data input (1) is a single matrix or two matrices for cross-omics mode. Pairwise association (2) is then calculated between all molecules in the preprocessed matrix using the specified method. Evaluation (3) is calculated against a benchmark (when provided) and confidence is calculated for each prediction. Output is a ranked edge list with an association score and confidence (if calculated).

### 2.3 VaLPAS Autoencoder Architecture

The VaLPAS autoencoder is a novel architecture designed specifically for omics data where the number of features (p) typically far exceeds the number of conditions (n). Unlike standard autoencoders that operate on individual samples, this architecture simultaneously learns embeddings along both the omics feature (protein/gene/metabolite/etc.) and condition (sample) dimensions of the data matrix.

#### 2.3.1 Architecture Design

Given an input data matrix where p = number of features and n = number of conditions, two encoders encoding omics features and samples learn low-dimensional embeddings for each feature and sample respectively. The feature encoder by encoding the features expression profile across all conditions and the sample encoder by encoding the abundance levels of all features in that condition. The encoders are applied independently to each of the p features and n samples, producing an omics feature embedding matrix and sample embedding matrix. Decoding then occurs through a dual decoder and two parallel pathways where the omics feature decoder maps each feature embedding back to the sample space and vice versa. The final reconstruction is then a learned weighted combination of both decoder outputs with a learnable scalar parameter (α; initialized at 0.5) passed through a sigmoid function.

#### 2.3.2 Training Procedure

Training employs a denoising objective inspired by masked language modeling. At each training step a binary mask is randomly generated for 15% of the matrix entries, setting these to 0. This procedure eliminates the explicit need for train/test splits, as the masking procedure implicitly provides this at the input value level. The model then reconstructs the full matrix learning over 200 epochs (user specified) with the Adam optimizer (learning rate 1e-3, weight decay 1e-5) and a mean squared error (MSE) loss function.

This forces the model to learn feature co-expression patterns rather than memorizing individual values, providing implicit regularization that is particularly beneficial in the low-sample regime. The major limitation of this is approach is that the learned model is only applicable to the input data matrix and cannot be easily generalized to another input data set. However, this is in line with the statistical methods included in VaLPAS which have the same limitation.

#### 2.3.3 Similarity Extraction

After training, functional associations between features are derived from the learned feature embeddings using cosine similarity and converting the resulting similarity matrix into a set of edges sorted in descending order. The cosine similarity in embedding space captures non-linear co-expression relationships that linear correlations miss. Features with similar embedding vectors are inferred to share functional relationships. Results were tested through benchmarking (see Methods 2.4). As previously mentioned, this cosine similarity should not be confused with the cosine similarity based statistical analysis that does not utilize the autoencoder procedure.

#### 2.3.4 Cross-omics extension

For cross-omics analysis (e.g., protein-metabolite associations) using the autoencoder-based association method, VaLPAS concatenates the two measurement matrices along the feature axis after aligning on shared conditions. The autoencoder then operates on the combined matrix, learning embeddings that capture both within-omics and between-omics relationships. Cross-omics associations appear as cosine similarities between embedding vectors from different omics types.

### 2.4 Benchmarking

#### 2.4.1 Gold standard known relationships

KEGG (Kanehisa, et al., 2025) reactions, modules, and orthology mapping were retrieved using KEGG’s REST-style API using custom scripts (available in the accompanying paper repository). Co-membership in KEGG modules, small subnetworks of functionally related enzymes and metabolites that work together to accomplish a metabolic process, was used as one evaluation dataset. This provides continuity across organisms, as all organisms are mapped to the same set of metabolically important modules, and a way to assess the cross-omic predictions for metabolites. Additionally, relationships were retrieved from STRING (v12.0) using the STRING API (Szklarczyk, et al., 2025), specifically: experimental, database, textmining, fusion (domains that are sometimes fused together in one protein), and neighborhood information. Co-expression relationships were excluded, as this would result in circular evaluation. Finally, STRING interactions were also combined into an overall set (‘combined’).

The TIGRfams set of hidden Markov models is largely comprised of whole-protein (as opposed to functional domain) models constructed to classify proteins into families in which members share function as well as sequence similarity. The hmmer v3.3 software suite (https://hmmer.org) (Eddy, 2011) was used to search TIGRfams v15.0 models (Haft, et al., 2013; Li, et al., 2021) against protein data sets.

#### 2.4.2 Performance metrics

We define true positives (TP) as being those predictions by VaLPAS at the specified score threshold or prediction rank depending on purpose, that are present in the gold standard benchmark set being considered. True negatives (TN) are those predictions which match a permutation of all individual nodes (gene, protein, metabolite) that are in the TP set, but that are not TPs themselves.

For overall classification performance, area under the receiver-operator characteristic curve (AUROC) was calculated using the included functions in scikit-learn.

Confidence of predictions was calculated as the positive predictive value (PPV) for the top n predictions, where n is the rank of the predicted association. PPV is calculated as:

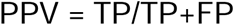

Where VaLPAS considers all combinations of input nodes that are not in the set of known associations (positives; P) to be not true edges (negatives; N).

### 2.5 Implementation and availability

VaLPAS is implemented in Python and can be executed from the command line or using an API from workflow managers or Jupyter notebooks. VaLPAS relies on packages including bspline-mutual-information, numpy, networkx, pandas, and scikit-learn. A Jupyter notebook and an accompanying dataset is included providing examples of use and output results.

VaLPAS code, documentation, and simple examples are available at http://github.com/PNNL-Predictive-Phenomics/valpas. Additionally, a complete set of hypotheses, test scripts, figure generation scripts, and data to recreate the results in this paper is available at http://github.com/PNNL-Predictive-Phenomics/valpas-paper. This repository is a Scientific Method using AI Research Template (SMAIRT) project. SMAIRT is available at https://github.com/PNNL-Predictive-Phenomics/smairt-template. Though a portion of the VaLPAS code and the code used to generate results of this paper were generated using AI (Claude Opus 4.7), the authors have carefully validated and are responsible for all code and results in the paper.

## 3. Results

### 3.1 Benchmarking Pearson and autoencoder for prediction of known protein relationships

We initially compared the ability of Pearson correlation and the VaLPAS autoencoder to determine biologically accurate functional associations using bacterial transcriptomics datasets from four different species (*E. coli*, *P. putida*, *S. enterica*, *S. elongatus*, see Table 1), which have large numbers of samples. To assess performance, we examined whether each method could infer associations between genes that appear in the same known STRING relationship (see Methods 2.4). We hypothesized that increasing the number of samples considered would improve performance due to the increase in signal provided, and that the more sophisticated autoencoder method would outperform simple correlation. The overall performance of these approaches as evaluated by AUROC (Figure 2) supported our hypothesis, showing increasing performance as the number of samples increased. The overall performance was significantly above random (0.5), but relatively modest, and showed variable results across the four species tested. The autoencoder outperformed Pearson correlation in performance, supporting our hypothesis. Benchmarking VaLPAS against co-membership in KEGG modules (Figure S1) showed the same trend albeit not as clear as benchmarking against STRING. In the case of E.coli Pearson correlation performs as good if not slightly better than the autoencoder for a sample size of 200. This likely reflects differences in the kind of relationships the two benchmarks represent (see Discussion). Additionally, we observed that using individual STRING evidence types (e.g. neighborhood, experimental, etc.) as opposed to the combined evidence type (see Methods 2.4.1) resulted in different levels of performance as measured by AUROC, with Neighborhood only evaluation generally showing the best performance (Figure S2). Finally, the effect of noise on performance was examined for the *P. putida* dataset by randomly permuting the transcript abundance of individual samples from no permutation to 100% permutation. Both methods (Pearson and VaLPAS autoencoder) were fairly robust to this noise, maintaining over 50% performance at 75% sample scrambling (Figure S3).

**Figure 2.**
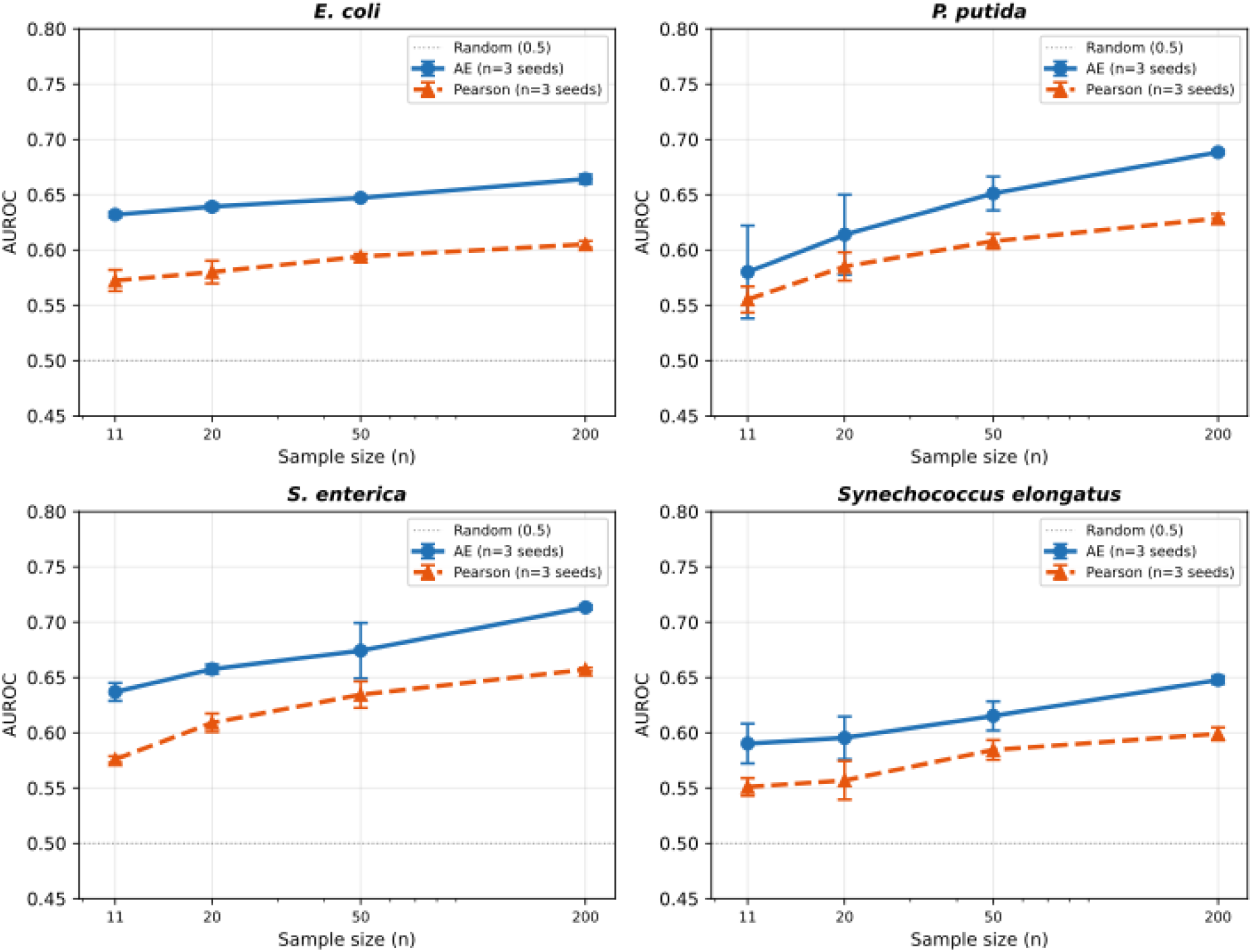
Comparison of VaLPAS functional association across bacterial transcriptomics datasets. Associations between genes using transcriptomics data for each strain (labels) was calculated using Pearson correlation (dotted lines) or the VaLPAS autoencoder (solid lines) for different sample sizes (X axis). Error bars show standard deviation of performance over three different random seeds. Performance was assessed as AUROC of the method against the STRING benchmark.

The modest overall performance of VaLPAS at determining relationships between genes is not surprising given that the transcriptomics datasets are limited, and not all genes will display the degree of variation needed for robust association. This observation highlights that the value of the approach is its ability to screen for interesting predictions, rather than a complete annotation tool itself. Therefore, we next evaluated how the association score for Pearson correlation or the autoencoder relates to the confidence (evaluated as PPV) for each of the bacterial transcriptomics datasets. For at least one dataset (*E. coli*), the autoencoder was found to have compressed score values, so we evaluated the PPV of the prediction against the rank of that prediction as provided by the score, effectively normalizing score ranges between approaches and datasets. This analysis applied across bacterial species using 200 samples each, reveals that there is a direct relation between the two measures, specifically that higher association scores (lower ranks) directly translate into higher confidence (Figure 3). These results are notable in that they show that although the Pearson correlation and autoencoder curves are somewhat different, both provide high confidence predictions at the higher ranked predictions. Each dataset differs slightly, however, in terms of relative performance Pearson correlation shows an advantage for *E. coli* while the VaLPAS autoencoder performs either equally (*S. typhimurium*) or better (*P. putida* and *S. elongatus*). For all but the *S. elongatus* data, both methods are able to provide more than 1000 predictions at confidence levels of 50% or higher (Figure 3, points).

**Figure 3.**
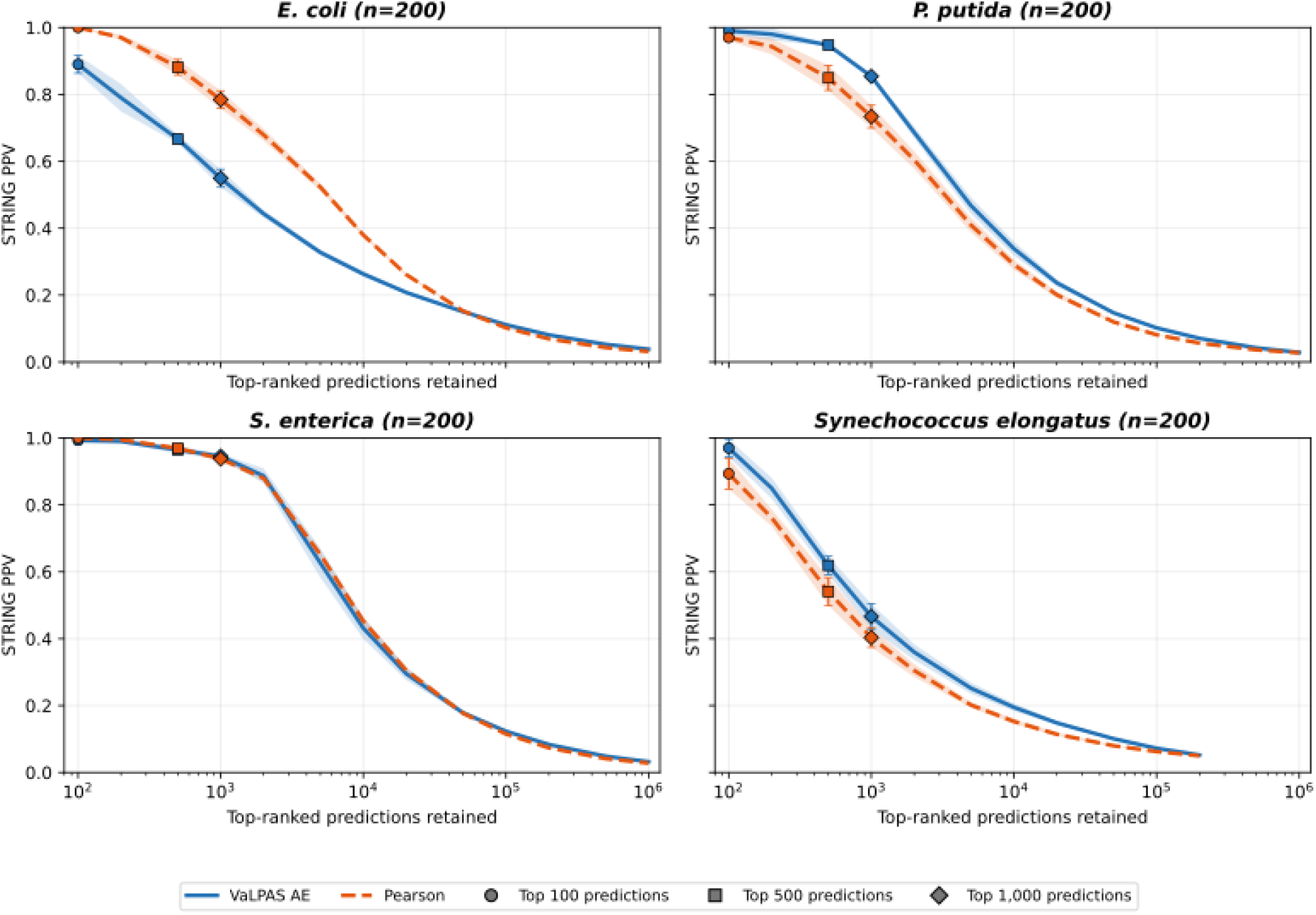
Confidence calibration for bacterial transcriptomics. The PPV evaluated using STRING is shown (Y axis) versus the prediction rank threshold by score (X axis; log scale) for VaLPAS autoencoder (solid lines) or Pearson correlation (dotted lines). Points mark where on the curve where 100, 500, and 1000 top predictions fall. Shaded regions on each plot show standard deviation from three random seeds, and bars show standard deviation at noted points.

To evaluate the confidence of predictions that could be obtained with a smaller set of transcriptomics data, as might be more feasible to produce in a single experimental study, the same analysis was performed for *P. putida* using a sample size of 20. The result of this analysis (Figure S4) shows that VaLPAS can provide high-quality (confidence > 70%) for 100 predictions, and reasonable quality (>30%) for up to 1000 predictions.

As bacterial regulation is known to be simpler than eukaryotic regulation, we were interested in testing whether similar results would be seen when making predictions with eukaryotic omics datasets. We compared these approaches in transcriptomics datasets from the fungi *Saccharomyces cerevisae*, using the same benchmarking approach. These results (Figure 4) show that the VaLPAS autoencoder is significantly better at lower sample numbers down to 11, but that Pearson reaches or exceeds the autoencoder performance only with a larger number (200) of samples which are available for *S. cerevisiae*. Similar results were obtained using KEGG as a benchmark (Figure S5).

**Figure 4.**
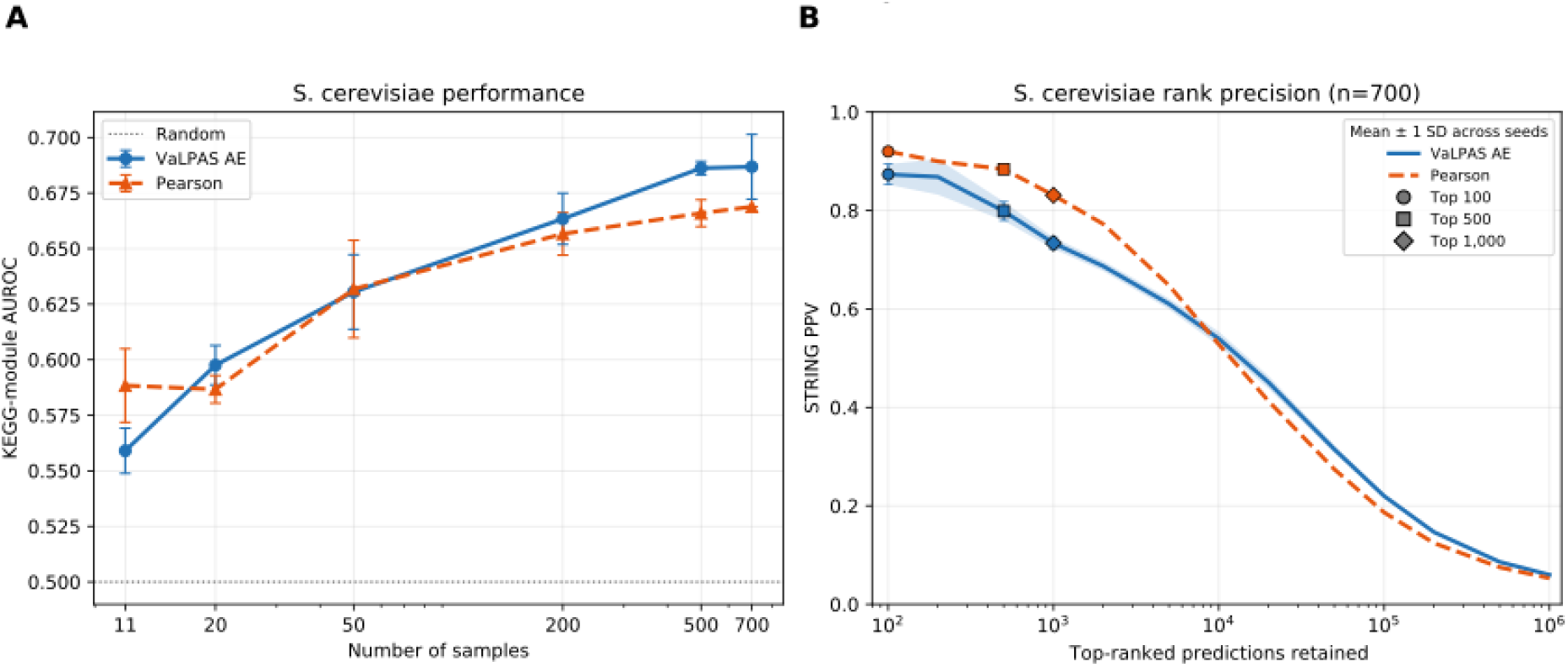
VaLPAS can provide high-confidence functional associations using fungal transcriptomics. **A.** VaLPAS (autoencoder or Pearson association) performance as AUROC prediction (Y axis) on *S. cerevisiae* transcriptomics datasets of varying sizes (X axis) showing comparable overall performance as bacterial datasets. **B.** For the *S. cerevisiae* dataset at 200 samples the confidence calculated as PPV on known STRING relationships (Y axis) is shown versus the predictions ranked by score for VaLPAS Pearson or autoencoder associations (X axis) showing that high-confidence associations are provided at high score thresholds for both methods.

### 3.2 Characterization of dark genes in Pseudomonas putida using correlation and autoencoder approaches

Previously, Tavis and Hettich (Tavis and Hettich, 2024) have reported an integrative method for annotating dark genes in *P. putida* and reporting the ability to provide confident annotations for over 200 proteins using sequence, structure, co-abundance, and other information. Though these are compelling results, they do not explicitly test our central question: can omics data alone provide useful predictions of function for dark molecules? In fact, the previous study showed limited effectiveness for proteomics co-abundance using a compendium of three studies with approximately 30 conditions using a random forest method. The ability of correlation and our autoencoder methods to provide useful annotations for dark genes using VaLPAS was compared (Figure 5). To do so the full *P. putida* transcriptomic compendium was analyzed, comprising 321 samples. We applied either Pearson correlation or VaLPAS autoencoder to the complete dataset (see Methods), then evaluated the resulting similarity relationships against known relationships between genes represented in the STRING database, omitting ‘co-expressed’ relationships (see Methods 2.4). Dark genes were assessed as those genes that were not annotated with a KEGG orthology identifier and had no matches to TIGRFAMs families by HMMER (see Methods 2.4.1), which identified 1,510 dark genes with transcriptomics data (Supplementary Table 1). Confidence values (PPV) for score thresholds were taken from the results shown in Figure 3. The number of dark genes associated with a well-annotated partner at each confidence level was counted for both the Pearson correlation and autoencoder approach. The results show that VaLPAS can make confident predictions for many dark genes (139) at reasonable confidence levels above 70%, but that Pearson correlation provides a larger number of potential annotations for this application.

**Figure 5.**
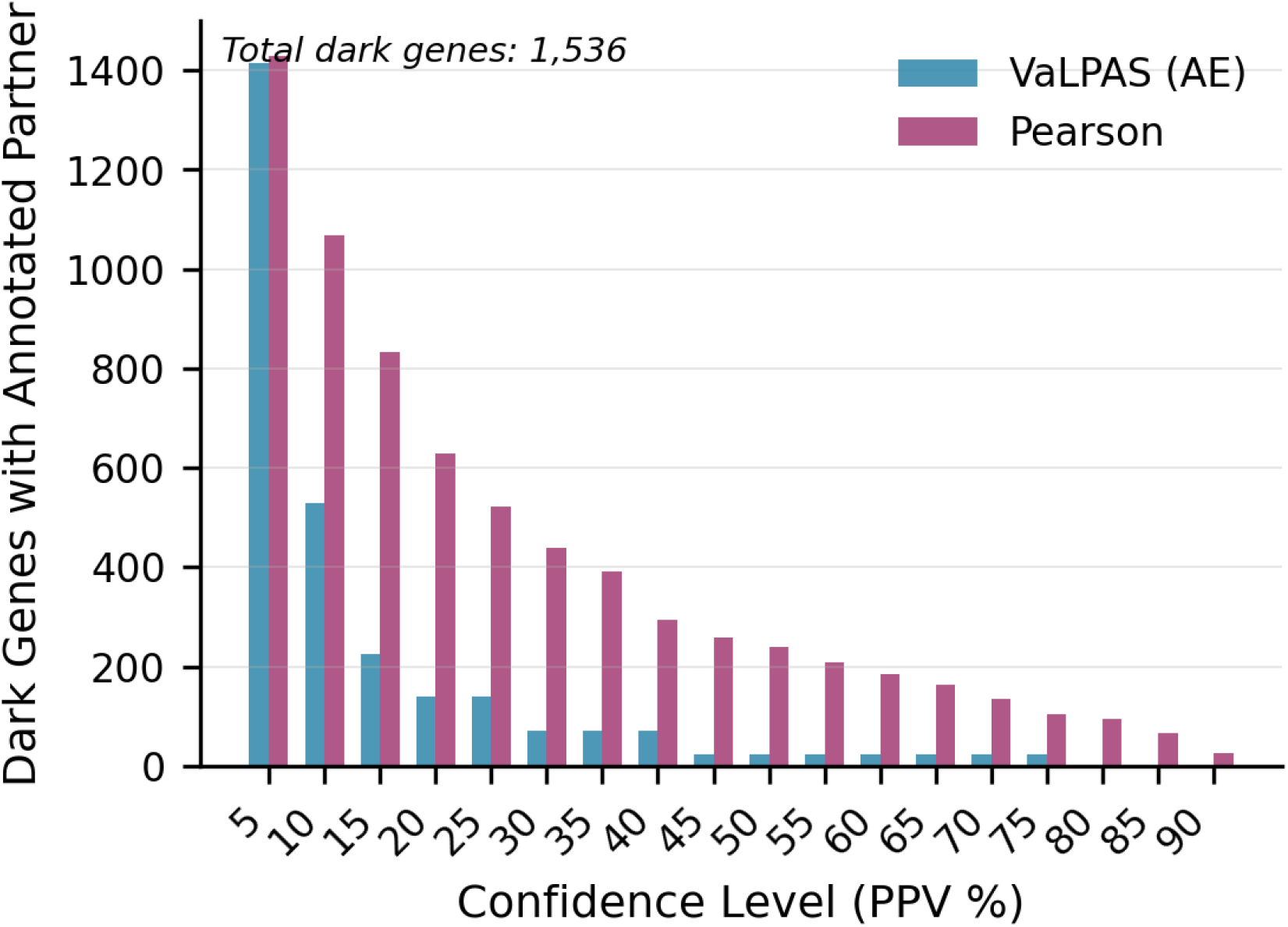
Dark gene association in *P. putida*. VaLPAS autoencoder or correlation was used to associate genes from 321 transcriptomics samples. Confidence was calculated for each prediction using STRING relationships and the number of dark genes, unannotated by KEGG or TIGRFams, that could be associated with an annotated partner at each confidence level is plotted for autoencoder (blue) or Pearson correlation (red).

We compared VaLPAS results with those of Tavis, et al. who reported a list of dark genes that their approach could annotate based on a guilt-by-association (GBA) method using several lines of evidence excluding their structure based results which are not comparable here. Their analysis yielded 9 dark genes annotated by GBA at the 90% confidence level that we have in our dark gene list, whereas VaLPAS could associate 26 at the corresponding level. However, these sets of predictions at the highest confidence do not overlap, highlighting the complementarity of approaches.

### 3.3 Characterization of dark metabolites in Lipomyces starkeyi using correlation and autoencoder approaches

The ability of Pearson correlation to annotate dark genes, where gene-gene relationships are expected to be near linear in nature, raised the hypothesis that the autoencoder approach would be better suited to cross-omics associations, such as those between enzymes and metabolites. We tested this hypothesis using our recently published multi-omics dataset from the oleaginous yeast, *Lipomyces starkeyi* in which matched proteomics and metabolomics data was acquired for samples grown under different carbon and nitrogen culture conditions (Monteiro, et al., 2026). Dark metabolites were determined as metabolic peaks identified by mass spectrometry and consistently observed across samples, but where no molecular identity could be assigned by the processing pipeline used in the paper. Analogous to how we identified dark genes, dark metabolite predictions were assigned confidence scores based on known KEGG relationships in pathways (Figure S6), and numbers reported if the dark metabolite was associated with a known protein at the indicated confidence level (Figure 6). In this case Pearson correlation was unable to provide predictions above 20% confidence, whereas the VaLPAS autoencoder approach associated 176 dark metabolites with proteins of known function at a confidence of 50% or higher (Supplementary Table 2).

**Figure 6.**
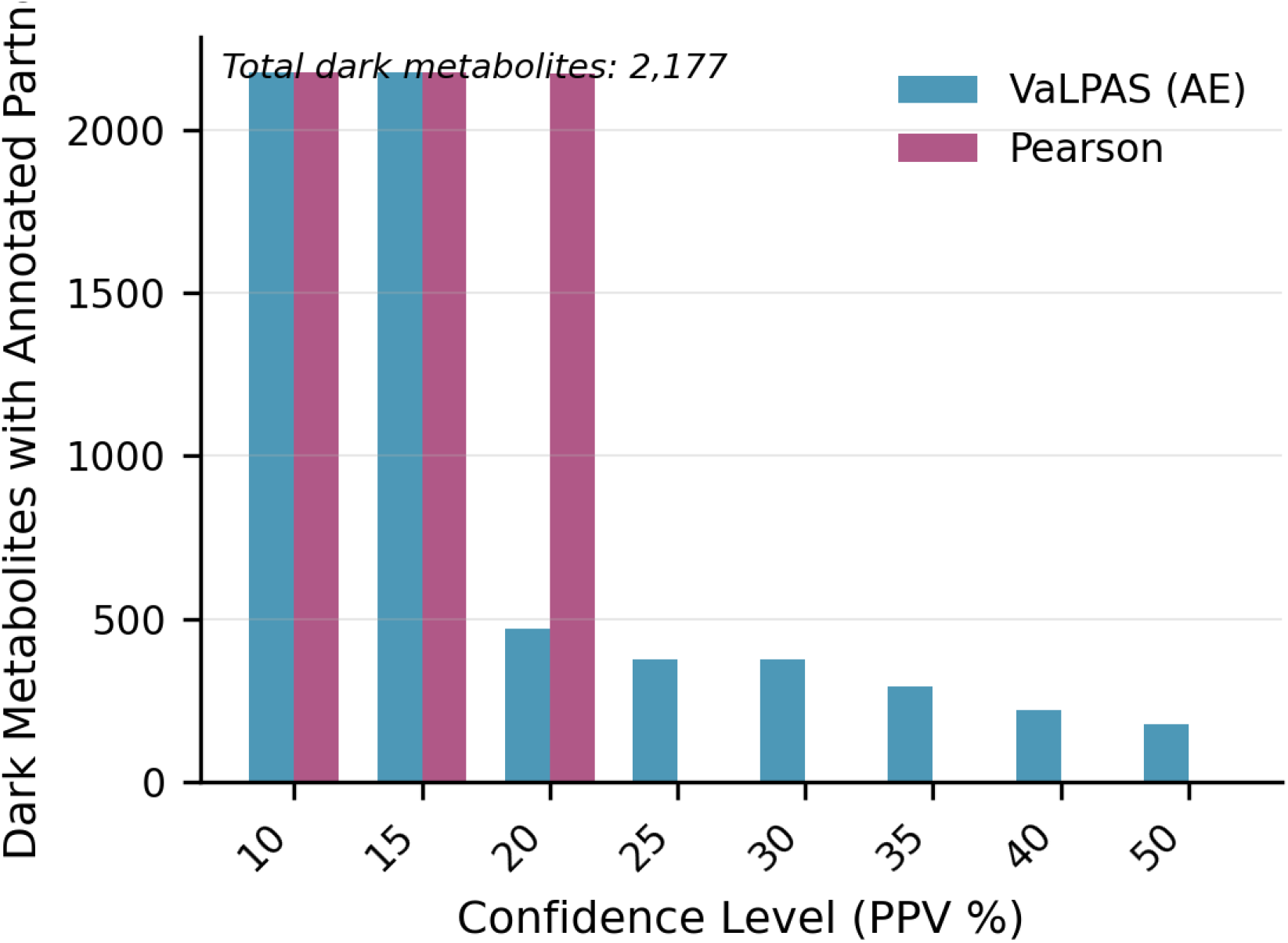
Dark metabolite association in *L. starkeyi*. VaLPAS was used to associate proteomics and metabolomics and confidence was calculated using KEGG pathway membership. The number of unknown metabolites (metabolic peaks with no assigned molecular identity) is shown plotted against the confidence threshold.

## 4. Discussion

To examine the effectiveness of association-based methods to enable annotation of dark molecules using omic and multi-omic datasets we have developed VaLPAS, a framework for calculation of association using omics data. VaLPAS includes standard statistical association methods such as correlation and mutual information as well as introducing a novel bi-directional autoencoder method for learning patterns from the input omics matrix. The framework has been designed with the specific intent to elucidate functional relationships between features within and across different omics modalities (e.g. between proteins and metabolites) and should not be confused with annotation tools. As such VaLPAS serves as an additional line of functional relationship evidence akin to individual protein interaction evidence in STRING, while not being limited to protein-protein relationships. Furthermore, in conjunction with preexisting evidence such as KEGG pathways, VaLPAS is able to identify co-membership of not just proteins, but other omics modalities associated with proteins inside of a given pathway. While not being strictly feature annotation, we argue it can elucidate functional information of previously unknown features.

As previously described and utilized (Wang, et al., 2017; Wolfe, et al., 2005), guilt-by-association methods like VaLPAS are most useful as by allowing insight into function of sets of proteins that are not amenable to traditional sequence-based methods. In our study we evaluated this directly by categorizing a set of dark genes, those that do not have a KEGG orthology assignment and do not have hits in the TIGRFAMs bacterial family database. In *Pseudomonas putida* we show that VaLPAS can identify partners with annotations for 130 dark genes at better than 75% confidence. Comparison at the highest confidence shows that VaLPAS identifies many more dark gene partners than did the method by Tavis, et al. It is important to note that this is not a direct comparison because each approach used very different sources and amounts of data, as well as different association methods, but it highlights that VaLPAS is a useful tool for characterization of dark genes in bacteria. This analysis showed that VaLPAS was able to provide annotations for genes that were not amenable to traditional and highly sensitive sequence similarity methods such as HMMER.

There are other approaches that utilize information beyond sequence for prediction of protein annotation. The S2F approach uses sequence similarity combined with network analysis derived from the STRING database (https://paccanarolab.org/s2f/). GeneMania combines integration of different networks with label propagation to provide enhanced annotations (Mostafavi, et al., 2008). NetGO uses sequence-based similarity combined with network-based annotation using existing knowledge graphs (Wang, et al., 2023). Since the purpose of VaLPAS is to utilize only associations derived from omics datasets to predict molecular function relations, we see these other tools as representing complementary methods for protein annotation. We show in this paper an additional distinction that VaLPAS is able to provide functional roles for unknown metabolites at reasonable confidence given appropriate multi-omics datasets, something that is not possible with other tools.

Given the results presented here we believe that VaLPAS is a significant step forward in the use of omics data to provide predictions of function for dark molecules. However, our results highlight several limitations in this approach. The first is that VaLPAS requires experiments with many different experimental conditions to achieve its full performance and thus may be best suited to analysis of compendia such as those available for transcriptomics that we have utilized here. VaLPAS provides the ability to look across omics data, specifically at dark metabolites, but requires experiments with multiple conditions for which matched proteomics and metabolomics data are available. Though the number of these kinds of datasets is growing (Schorn, et al., 2021) this requirement is limiting. We are currently investigating methods to approach this problem that do not require this strictly sample-matched data, which would make it more accessible to the community. Prediction of dark metabolites is a significant step forward, providing a novel method to address this gap, but the results we present are based on a limited dataset, show modest confidence at the highest score thresholds, and provide general hypotheses for metabolite identity at the pathway level. We believe that combining this approach with established chemoinformatics approaches, as prior probabilities to guide searches, will be a useful future direction for investigation. Finally, we note that the results presented here show variability from dataset to dataset, indicating that some caution should be taken, especially when using VaLPAS for less well-studied organisms (see *A. baumanni* results in Figures 2 and 3). If provided a list of known functional relationships, however, the VaLPAS framework will automatically calculate confidence levels for predictions, allowing users to evaluate predictions critically.

## 5. Conclusion

Our current analysis spanning four bacterial species and three fungal species, shows that VaLPAS is able to provide high confidence functional associations for a significant number of genes, even with relatively modest numbers of transcriptomics samples. Our transcriptomics analysis shows that Pearson correlation performs surprisingly well, but that the novel VaLPAS autoencoder architecture outperforms correlation for most use cases. Additionally, our cross-omics analysis shows that the VaLPAS autoencoder can learn relationships between protein and metabolites, and that this approach provides the only method we are aware of to link unidentified metabolic peaks (putative novel metabolites) with known proteins and pathways. We note that these results are based on a small number of samples representing 36 samples from nine experimental conditions in a single study, and that we expect the performance of the method to improve as larger datasets are considered.

We believe that VaLPAS represents a significant step forward in elucidating dark functional matter, distinguishing itself from most existing functional annotation tools by being able to predict functional associations between various omics modalities. Importantly, VaLPAS enables linking uncharacterized small molecules with proteins and vice versa, and as such serves as a tool for generating hypotheses of functional interaction that can be validated with targeted experiments.

## Acknowledgements

We want to extend thanks to Sneha Couvillion and Jeff Czajka for vital feedback and scientific discussion in regard the interpretation multi-omic data.

Some portions of the VaLPAS code were generated using AI (Claude Sonnet 4.0) and then extensively tested and revised as needed by the authors to ensure accuracy.

## Author Contributions

Yannick Mahlich: Analysis, Methodology, Software – development, Writing – original draft & editing; Dylan H. Ross: Software – testing, Writing – review & editing; Lummy Monteiro: Software – testing, Writing – review & editing; Sutanay Choudhury: Writing – review & editing; William C. Nelson: Analysis, Writing – review & editing; Jason E. McDermott: Analysis, Conceptualization, Software – development, Supervision, Writing – original draft & editing

## Funding

Funding was provided from Pacific Northwest National Laboratory’s Predictive Phenomics Initiative conducted under the Laboratory Directed Research and Development Program. PNNL is a multiprogram national laboratory operated by Battelle for the U.S. Department of Energy under Contract No. DE-AC05-76RL01830

## Conflict of Interest

None

## Notes

### Competing Interest Statement

The authors have declared no competing interest.

### Summary of Updates

Update large portions of the text; authors updated; added supplementary figures and tables

