## Supplementary Figures for "Illuminating dark genes and dark metabolites: calibrated functional association across kingdoms with autoencoders and correlation"

Figure S1. Comparison of VaLPAS functional association across bacterial transcriptomics datasets using KEGG benchmark.

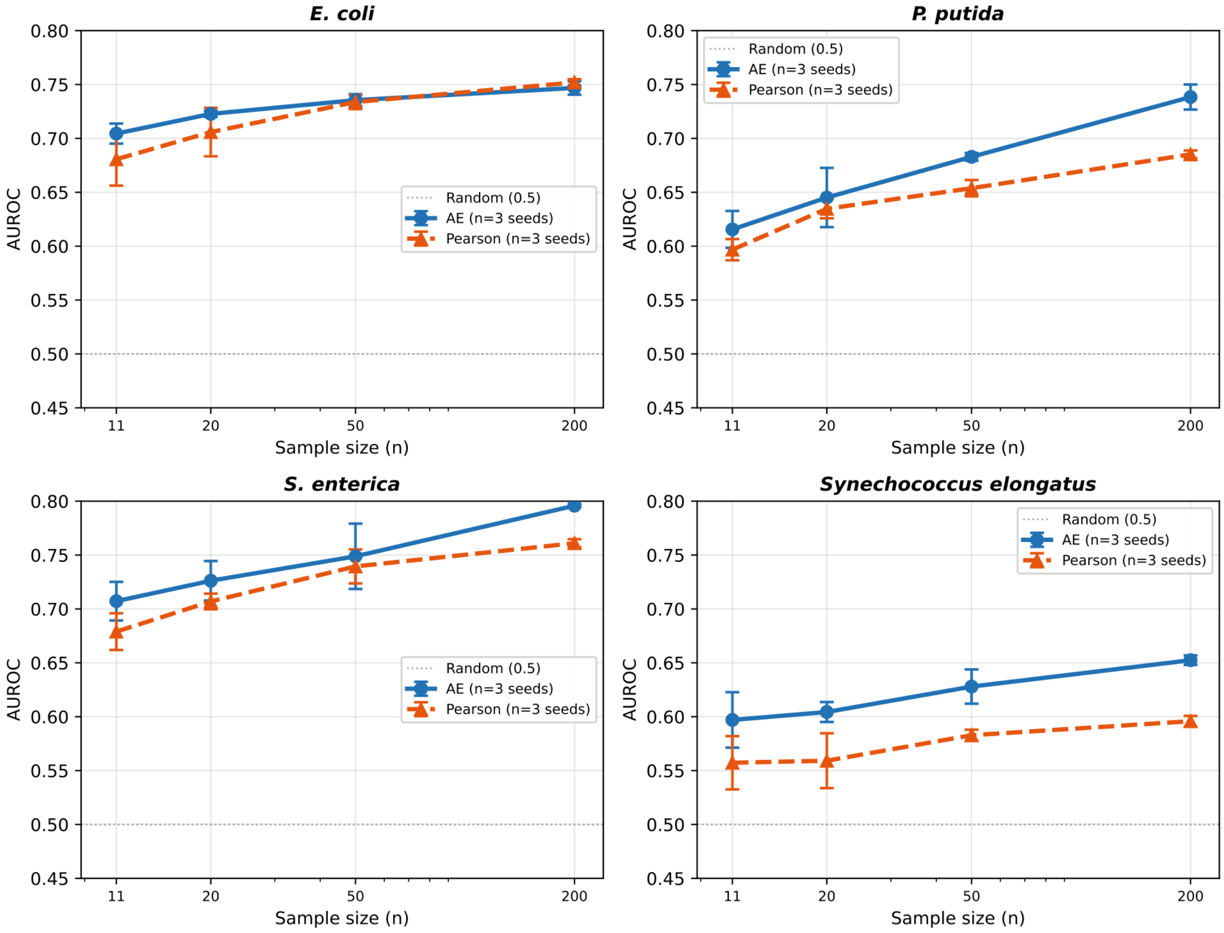

**Figure S1. Comparison of VaLPAS functional association across bacterial transcriptomics datasets using KEGG benchmark.** Associations between genes using transcriptomics data for each strain (labels) was calculated using Pearson correlation (dotted lines) or the VaLPAS autoencoder (solid lines) for different sample sizes (X axis). Error bars show standard deviation of performance over three different random seeds. Performance was assessed as AUROC of the method against the KEGG benchmark dataset.

Figure S2. Benchmarking associations against individual STRING evidence sources shows differences in performance.

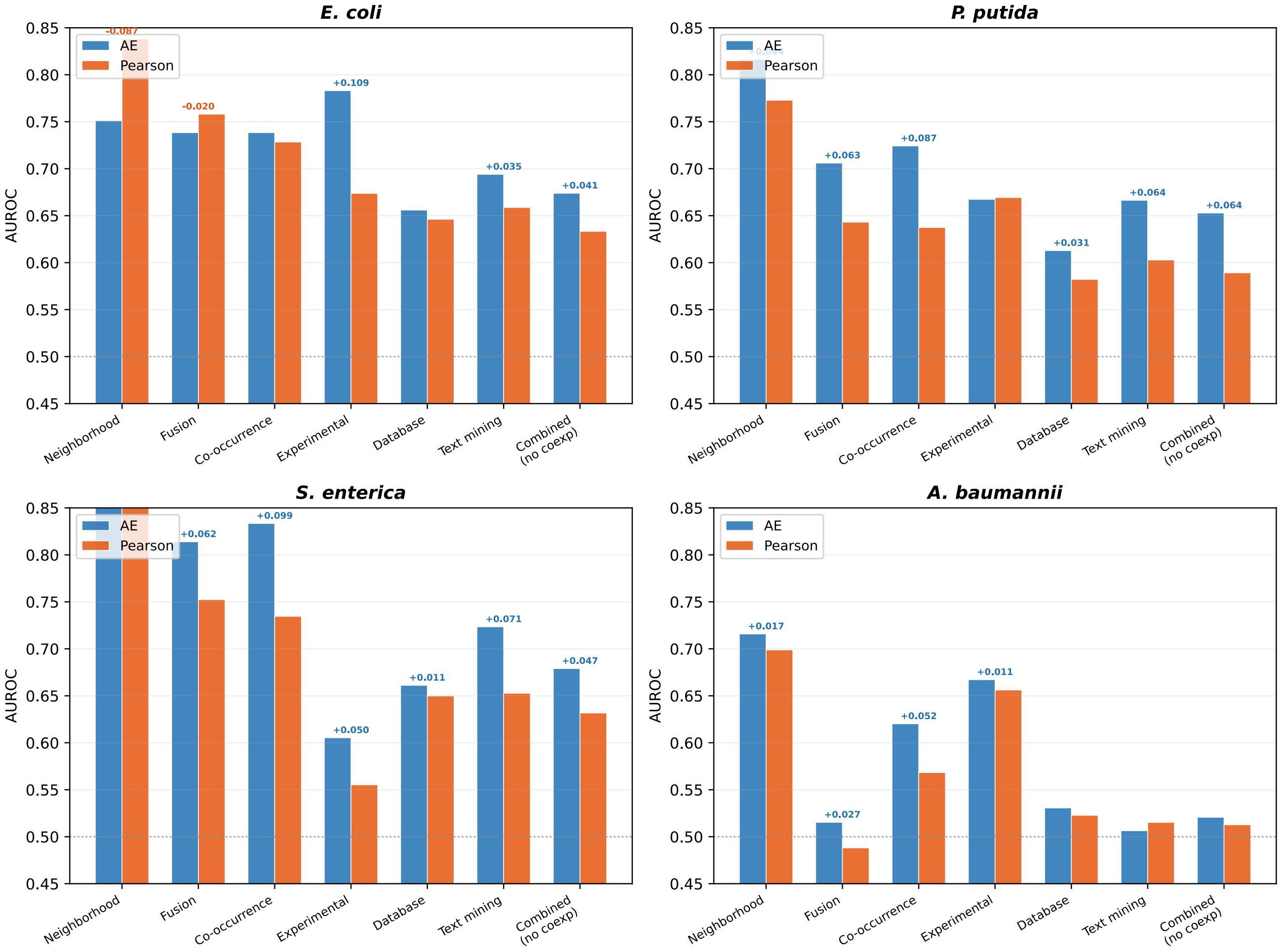

**Figure S2. Benchmarking associations against individual STRING evidence sources shows differences in performance.** For each bacterial dataset association scores were calculated using Pearson correlation (orange bars) and the autoencoder (blue bars) respectively. Benchmarking against individual STRING evidence sources (x-axis) shows different measured performance per source, with VaLPAS generally being able to recapture neighborhood derived associations in STRING the most reliable independent of association metric.

Figure S3. VaLPAS association robust to sample scrambling

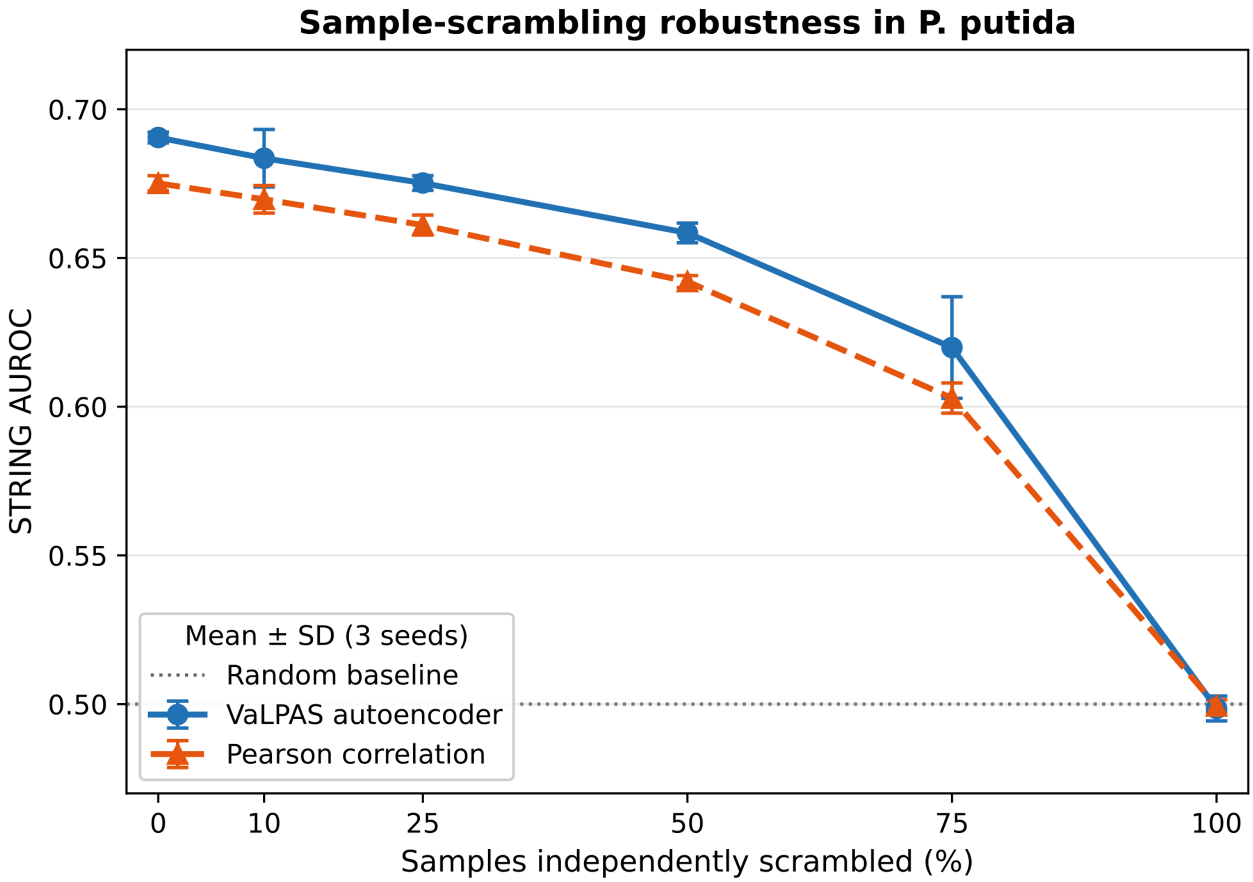

**Figure S3. VaLPAS association robust to sample scrambling.** Transcript abundances of individual samples in *P. putida* were randomly permutated from no permutation (x-Axis; 0%) to full (100% scrambling). AUROC was calculated for detected associations against STRING after abundance scrambling.

Figure S4. Confidence estimate in P. putida for small set of 20 samples.

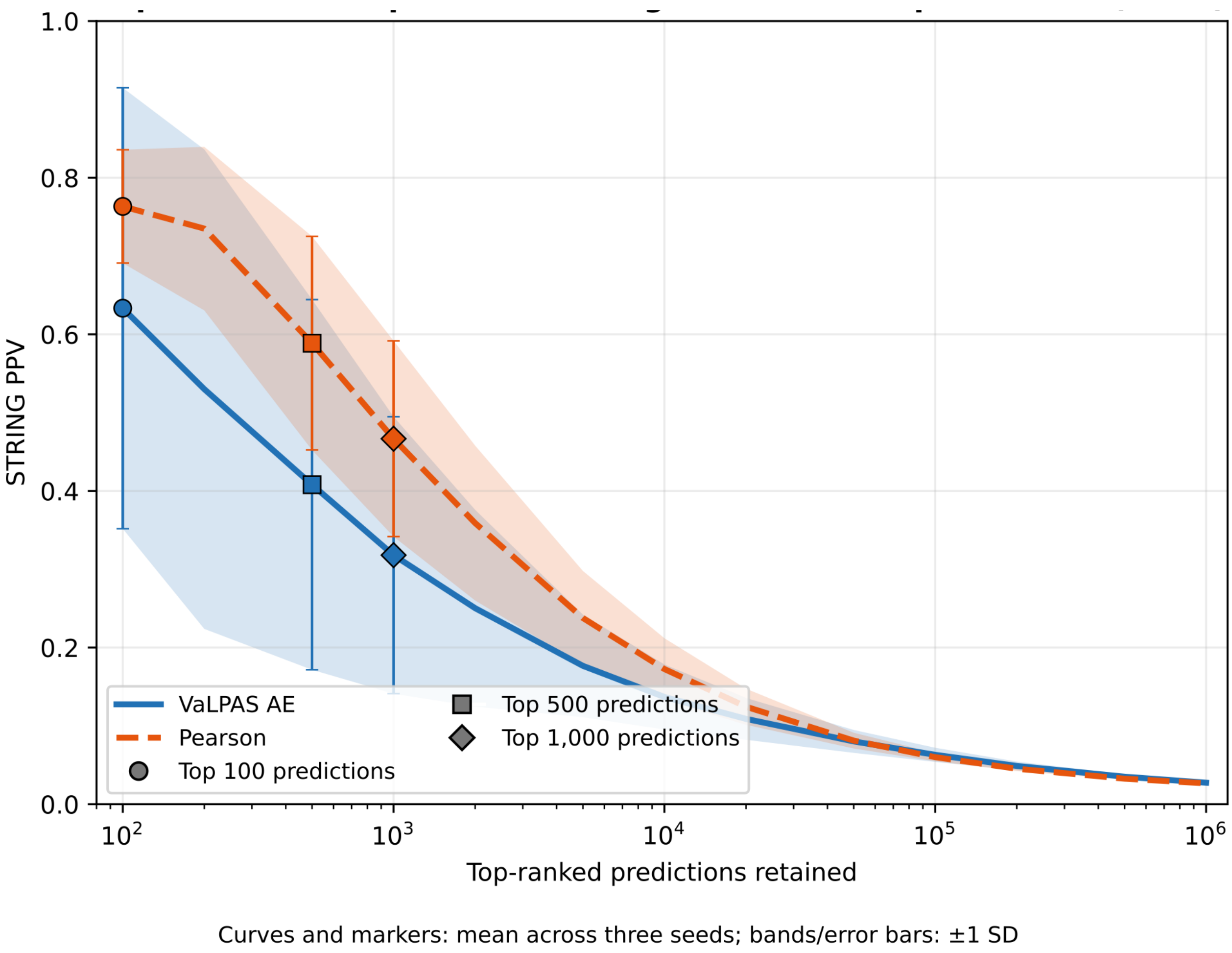

**Figure S4. Confidence estimate in P. putida for small set of 20 samples.** Overall performance is decreased in comparison to a larger set of samples (see Figure 3) but still reasonable. Additionally, Pearson performs better than the autoencoder in experiments with small sample sizes.

Figure S5. VaLPAS provided fungal transcriptomics functional associations benchmarked against KEGG modules

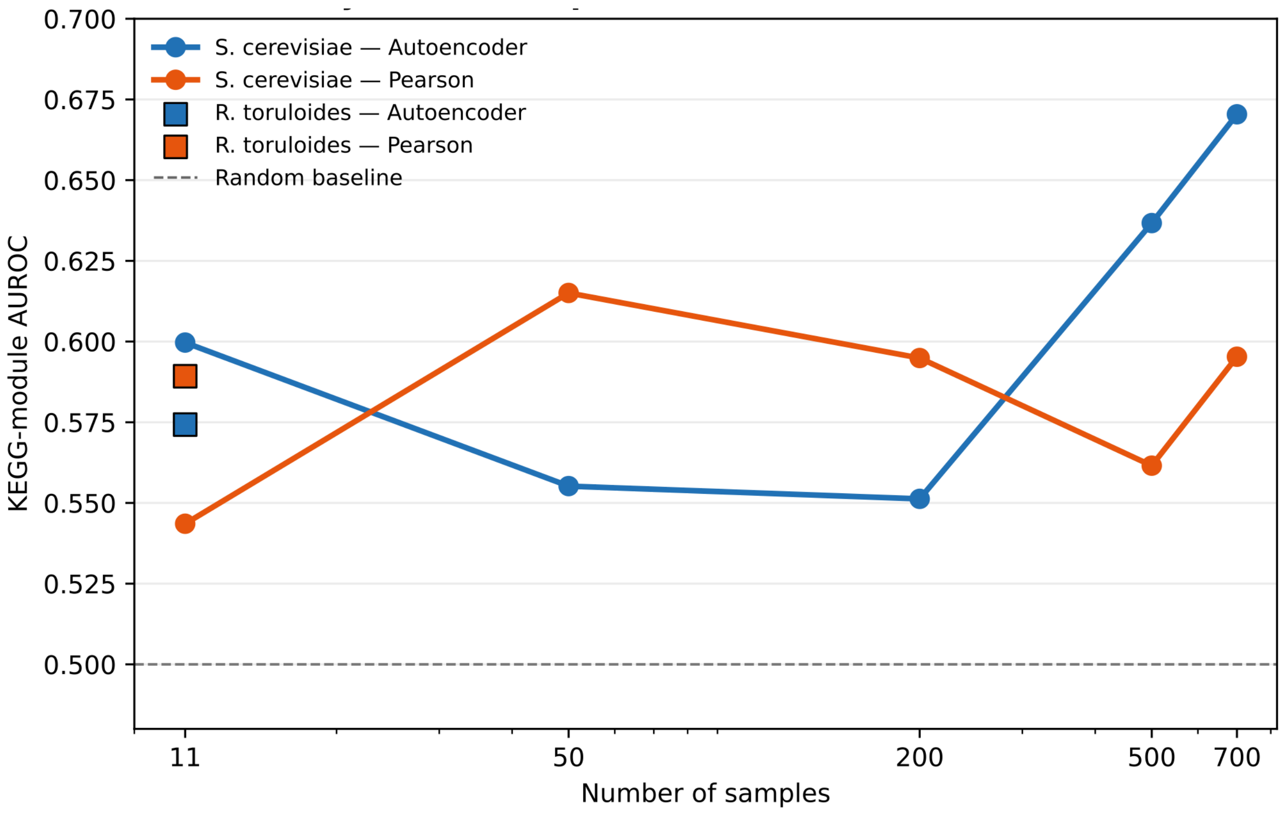

**Figure S5. VaLPAS provided fungal transcriptomics functional associations benchmarked against KEGG modules.** VaLPAS shows similar performance for both Pearson correlation and autoencoder based associations when benchmarked against co-membership in KEGG modules compared to STRING benchmarking (see Figure 4).

Figure S6. Cross-omics calibration for metabolite to protein associations

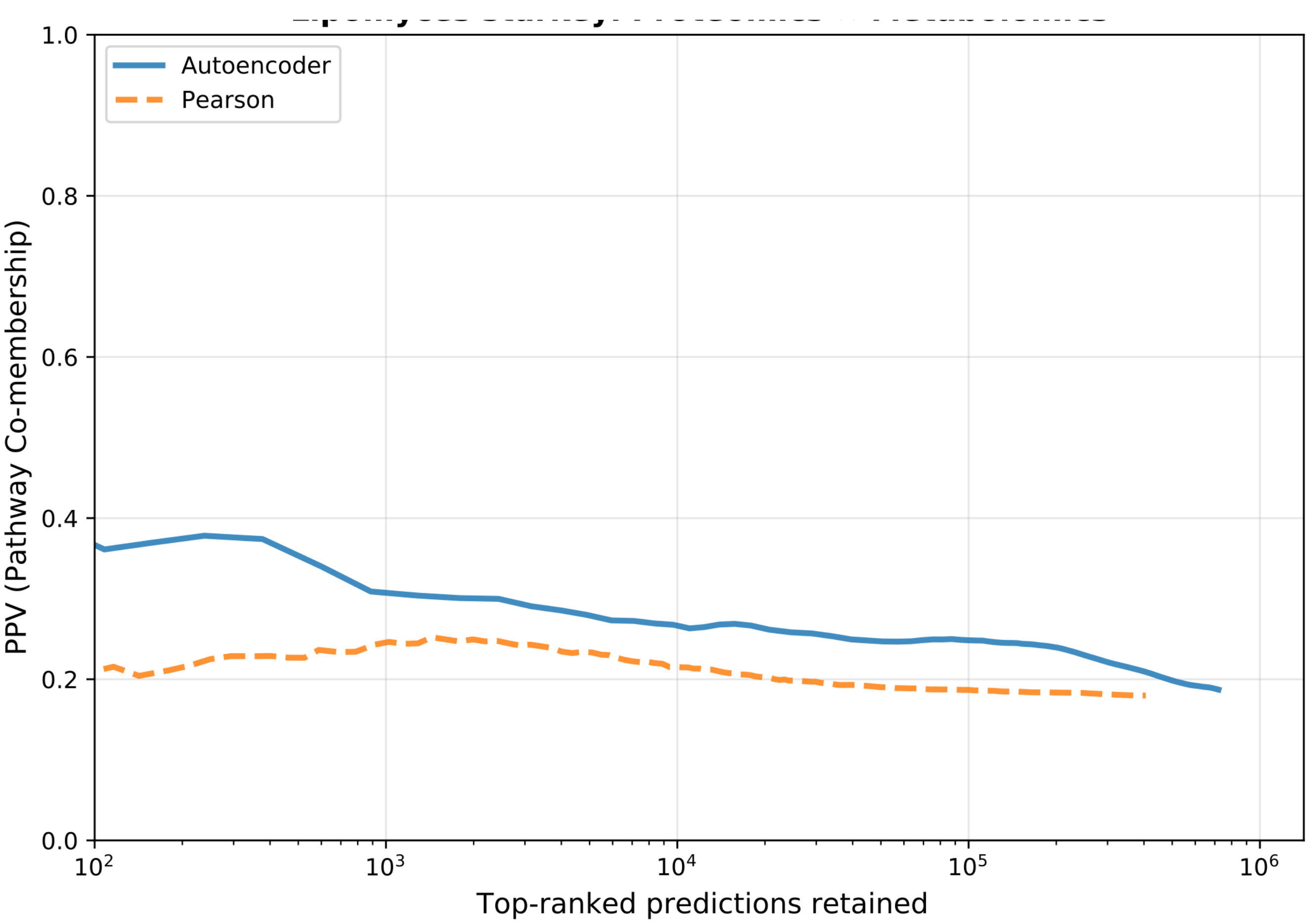

**Figure S6. Cross-omics calibration for metabolite to protein associations.** While overall confidence as evaluated by Positive Predictive Value benchmarked against KEEG pathway co-membership is relatively low, autoencoder based associations (blue) significantly outperform Pearson Correlation.
